# A Radically New Theory of how the Brain Represents and Computes with Probabilities

**DOI:** 10.1101/162941

**Authors:** Gerard (Rod) Rinkus

**Affiliations:** Neurithmic Systems, Newton, MA 02465 USA

**Keywords:** Sparse distributed representations, probabilistic population coding, cell assemblies, canonical cortical circuit/algorithm

## Abstract

Many believe that the brain implements probabilistic reasoning and that it represents information via some form of population (distributed) code. Most prior probabilistic population coding (PPC) theories share basic properties: 1) continuous-valued units; 2) fully/densely distributed codes; 3) graded synapses; 4) rate coding; 5) units have innate low-complexity, usually unimodal, tuning functions (TFs); and 6) units are intrinsically noisy and noise is generally considered harmful. I describe a radically different theory that assumes: 1) binary units; 2) sparse distributed codes (SDC); 3) *functionally* binary synapses; 4) a novel, *atemporal*, combinatorial spike code; 5) units initially have flat TFs (all weights zero); and 6) noise is a resource generated/used, normatively, to cause similar inputs to map to similar codes. The theory, Sparsey, was introduced 25+ years ago as: a) an explanation of the physical/computational relationship of episodic and semantic memory for the spatiotemporal (sequential) pattern domain; and b) a canonical, mesoscale cortical probabilistic circuit/algorithm possessing fixed-time, unsupervised, single-trial, non-optimization-based, unsupervised learning and fixed-time best-match (approximate) retrieval; but was not described as an alternative to PPC-type theories. Here, we show that: a) the active SDC in a Sparsey coding field (CF) simultaneously represents not only the likelihood of the single most likely input but the likelihoods of all hypotheses stored in the CF; and b) the whole explicit distribution can be sent, e.g., to a downstream CF, via a set of simultaneous single spikes from the neurons comprising the active SDC.

## 1 Introduction

It is widely believed that the brain implements some form of probabilistic reasoning to deal with uncertainty in the world [1], but exactly how the brain represents probabilities/likelihoods remains unknown [2, 3]. It is also widely agreed that the brain represents information with some form of distributed—a.k.a. population, cell-assembly, ensemble—code [see [4] for relevant review]. Several population-based probabilistic coding theories (PPC) have been put forth in recent decades, including those in which the state of all neurons comprising the population, i.e., the *population code*, is viewed as representing: a) the single most likely/probable input value/feature [5]; or b) the entire probability/likelihood distribution over features [6-11]. Despite their differences, these approaches share fundamental properties, a notable exception being the spike-based model of [11]. (1) Neural activation is continuous (graded). (2) *All* neurons in the coding field (CF) formally participate in the active code whether it represents a single hypothesis or a distribution over all hypotheses. Such a representation is referred to as a *fully distributed r*epresentation. (3) Synapse strength is continuous. (4) They are typically formulated in terms of rate-coding [12]. (5) They assume *a priori* that *tuning functions* (TFs) of the neurons are unimodal, e.g., bell-shaped, over any one dimension, and consequently do not explain how such TFs might naturally emerge, e.g., through a learning process. (6) Individual neurons are assumed to be intrinsically noisy, e.g., firing with Poisson variability, and noise is viewed primarily as a problem that to be dealt with, e.g., reducing noise correlation by averaging.

At a deeper level, it is clear that despite being framed as population models, they are really based on an underlying localist interpretation, specifically, that an individual neuron’s firing rate can be taken as a perhaps noisy estimate of the probability that a single preferred feature (or preferred value of a feature) is present in its receptive field [13], i.e., consistent with the “Neuron Doctrine”. While these models entail some method of combining the outputs of individual neurons, e.g., averaging, each neuron is viewed as providing its own individual, i.e., localist, estimate of the input feature. For example, this can be seen quite clearly in Fig. 1 of [9] wherein the first layer cells (sensory neurons) are unimodal and therefore can be viewed as detectors of the value at their modes (preferred stimulus) and the pooling cells are also in 1-to-1 correspondence with directions. This localist view is present in the other PPC models referenced above as well.

**Fig. 1.**
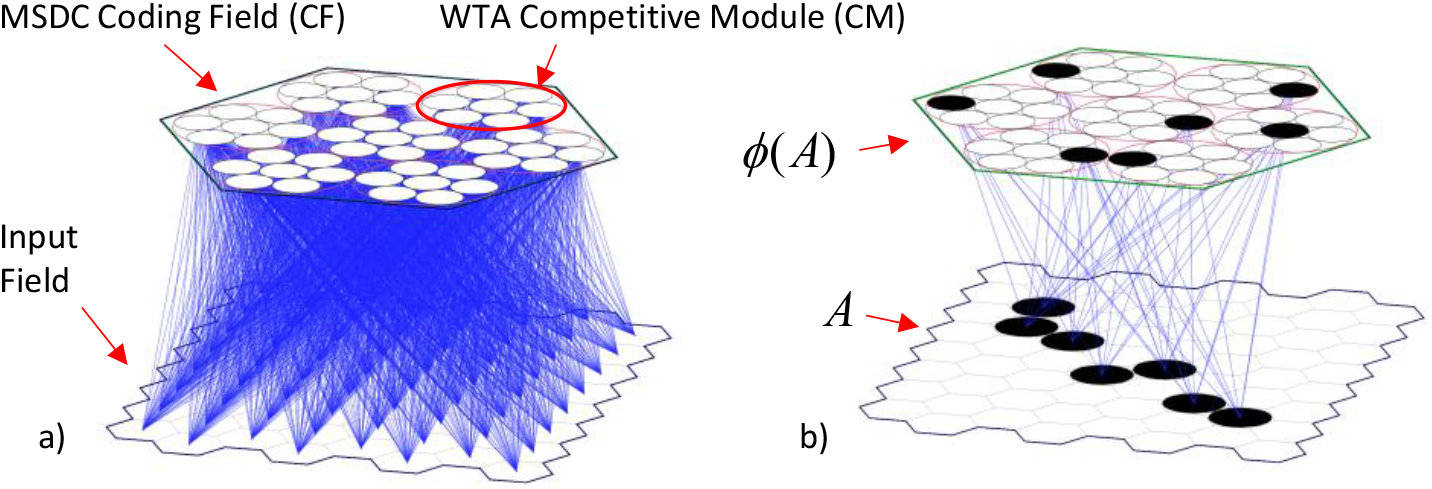
The *modular sparse distributed code* (MSDC) coding field (CF). See text.

However, there are compelling arguments against such localistically rooted conceptions. From an experimental standpoint, a growing body of research suggests that individual cell TFs are far more heterogeneous than classically conceived [14-21], also described as having “mixed selectivity” [22], and more generally, that sets (populations, ensembles) of cells, i.e., “cells assemblies” [23], constitute the fundamental representational units in the brain [24, 25]. And, the greater the fidelity with which the heterogeneity of TFs is modeled, the less neuronal response variation that needs to be attributed to noise, leading some to question the appropriateness of the traditional concept of a single neuron I/O function as an invariant TF plus noise [26]. From a computational standpoint, a clear limitation is that the maximum number of features/concepts, e.g., oriented edges, directions of movement, that can be stored in a localist coding field of *N* units is *N*. More importantly, as explained here, the efficiency, in terms of time and energy, with which features/concepts can be stored (learned) and retrieved/transmitted is far greater if items of information (memories, hypotheses) are represented with *sparse distributed codes* (SDCs) rather than localistically [27-29].

The theory described herein, Sparsey [27-29], constitutes a radically new way of representing and computing with probabilities, diverging from most existing PPC theories in many fundamental ways, including: (1) The representational units (principal cells) comprising a CF need only be *binary*. (2) Individual items (hypotheses) are represented by *fixed-size*, sparsely chosen subsets of the CF’s units, referred to as *modular sparse distributed codes* (MSDCs), or simply “codes” if unambiguous. (3) Decoding (read-out) not only of the most likely hypothesis but of the whole distribution, i.e., the likelihoods of *all* hypotheses stored in a CF, by downstream computations, requires only binary synapses. (4) The whole distribution, is sent via a wave of effectively simultaneous (i.e., occurring within some small window, e.g., at some phase of a local gamma cycle [30-33]) single spikes from the units comprising an active code to a down-stream (possibly recurrently to the source) CF. (5) The initial weights of all afferent synapses to a CF are zero, i.e., the TFs are completely flat. The classical, roughly uni-modal TFs [as would be revealed by low-complexity probes, e.g., oriented bars spanning a cell’s receptive field (RF), cf. [34]] emerge as a side-effect of the model’s single/few-trial learning process of storing MSDCs in superposition [35]. (6) Neurons are not assumed to be intrinsically noisy. However, the canonical, mesoscale (i.e., the cell assembly scale) circuit normatively uses noise as a resource during learning. Specifically, noise, presumably mediated by neuromodulators, e.g., ACh [36], NE [37], is explicitly injected into the code selection process to achieve the specific goal of (statistically, approximately) mapping more similar inputs to more similar MSDCs, where the similarity measure for MSDCs is intersection size. In this approach, patterns of correlation amongst principal cells are simply artifacts of this learning process.

### 2 The Model

Fig. 1a shows a small Sparsey model instance with an 8×8 binary units (pixel) input field that is fully connected, via binary weights (blue lines), all initially zero, to a *modular sparse distributed coding* (MSDC) coding field (CF). The CF consists of *Q* winner-take-all (WTA) *competitive modules* (CMs), each consisting of *K* binary neurons. Here, *Q*=7 and *K*=7. Thus, all codes have exactly *Q* active neurons and there are *K*^*Q*^ possible codes. We refer to the input field as the CF’s receptive field (RF). Fig. 1b shows a particular input A, which has been associated with a particular code, *ϕ*(A) (black units); here, blue lines indicate the bundle [cf. “Synapsemble”, [30]] of weights that would be increased from 0 to 1 to store this association (memory trace).

Fig. 2 illustrates MSDC’s key property that: ***whenever any <u>one</u> code is fully active in a CF, i*.*e***., ***all Q of its units are active, <u>all</u> codes stored in the CF will simultaneously be active (in superposition) in proportion to the sizes of their intersections with the single maximally active code***. Fig. 2 shows five hypothetical inputs, A-E, which have been learned, i.e., associated with codes, *ϕ*(A) - *ϕ*(E). These codes were manually chosen to illustrate the principle that *similar inputs* should map to *similar codes* (“SISC”). That is, inputs B to E have progressively smaller overlaps with A and therefore codes *ϕ*(B) to *ϕ*(E) have progressively smaller intersections with *ϕ*(A). Although these codes were manually chosen, Sparsey’s *Code Selection Algorithm* (CSA), described shortly, has been shown to statistically enforce SISC for both the spatial and spatiotemporal (sequential) input domains [27-29, 38, 39]: a simulation-backed example for the spatial domain is given in the Results section.

**Fig. 2.**
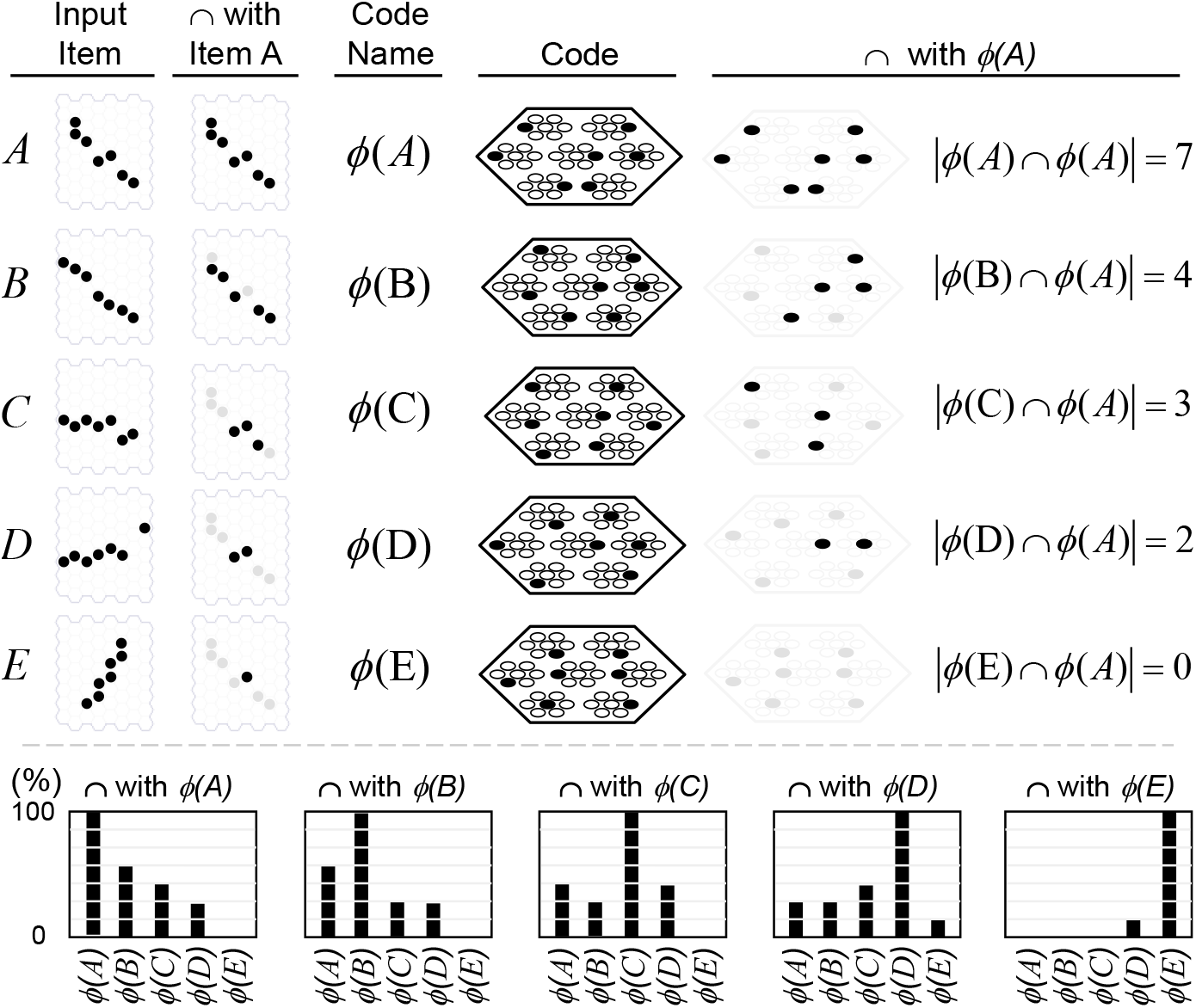
The probability/likelihood of a feature can be represented by the fraction of its code that is active. When *ϕ*(A) is fully active, the hypothesis that feature A is present can be considered maximally probable. Because the similarities of the other features to the most probable feature, A, correlate with their codes’ overlaps with *ϕ*(A), their probabilities/likelihoods are represented by the fractions of their codes that are active. In “⋂” columns, black units are those intersecting with the input A and with its code, *ϕ*(A); gray indicates non-intersecting units.

For input spaces for which it is plausible to assume that input similarity correlates with probability/likelihood, i.e., for vast regions of natural input spaces, the single active code can therefore also be viewed as a probability/likelihood distribution over all stored codes. This is shown in the lower part of Fig. 2. The leftmost panel at the bottom of Fig. 2 shows that when *ϕ*(A) is 100% active, the other codes are partially active in proportions that reflect the similarities of their corresponding inputs to A, and thus the probabilities/likelihoods of the inputs they represent. The remaining four panels show input similarity (probability/likelihood) approximately correlating with code overlap when each of the four other stored codes is maximally active.

### 2.1 The Learning Algorithm

A simplified version of the CSA, sufficient for this paper’s examples involving only purely spatial inputs, is given in Table 1 and we briefly summarize it here. [The full model handles spatiotemporal inputs, multiplicatively combining bottom-up, top-down, and horizontal (i.e., signals from codes active on the prior time step via recurrent synaptic matrices) inputs to a CF.] CSA Step 1 computes the raw input sums (*u*) for all *Q*×*K* cells comprising the coding field. In Step 2, these sums are normalized to *U* values, in [0,1]. All inputs are assumed to have the same number of active pixels, thus the normalizer, π_U_, can be constant. In Step 3, we find the max *U* in each CM and in Step 4, a measure of the familiarity of the input, *G*, is computed as the average max *U* across the *Q* CMs. In Steps 5 and 6, *G* is used to adjust the parameters of the nonlinear I/O transform in the same way for all of the CF’s units. In Step 7, each unit applies that “*U*-to-*μ”* transform, yielding an intermediate variable, *μ*, representing an unnormalized probability of the unit being chosen winner in its respective CM. Step 8 renormalizes *μ* values to total probabilities (*ρ*) of winning (within each CM) and Step 9 is the final draw from the *ρ* distribution in each CM resulting in the final code. *G*’s influence on the distributions can be summarized as follows.

**Table 1.**
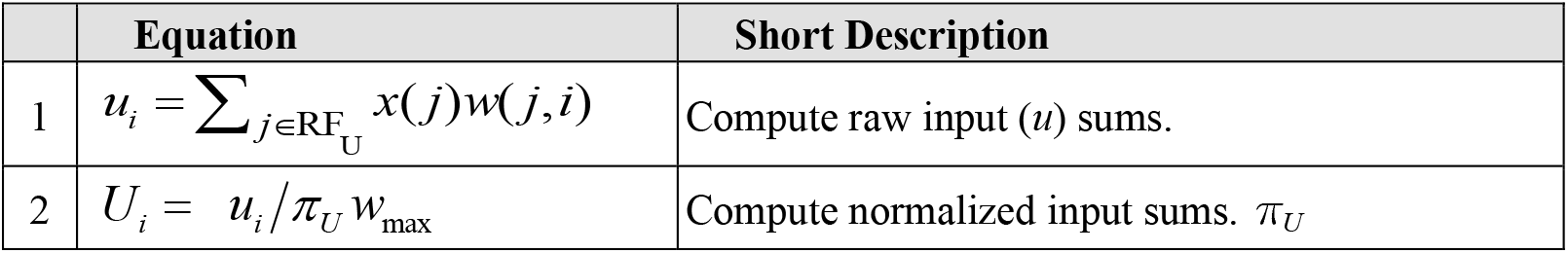
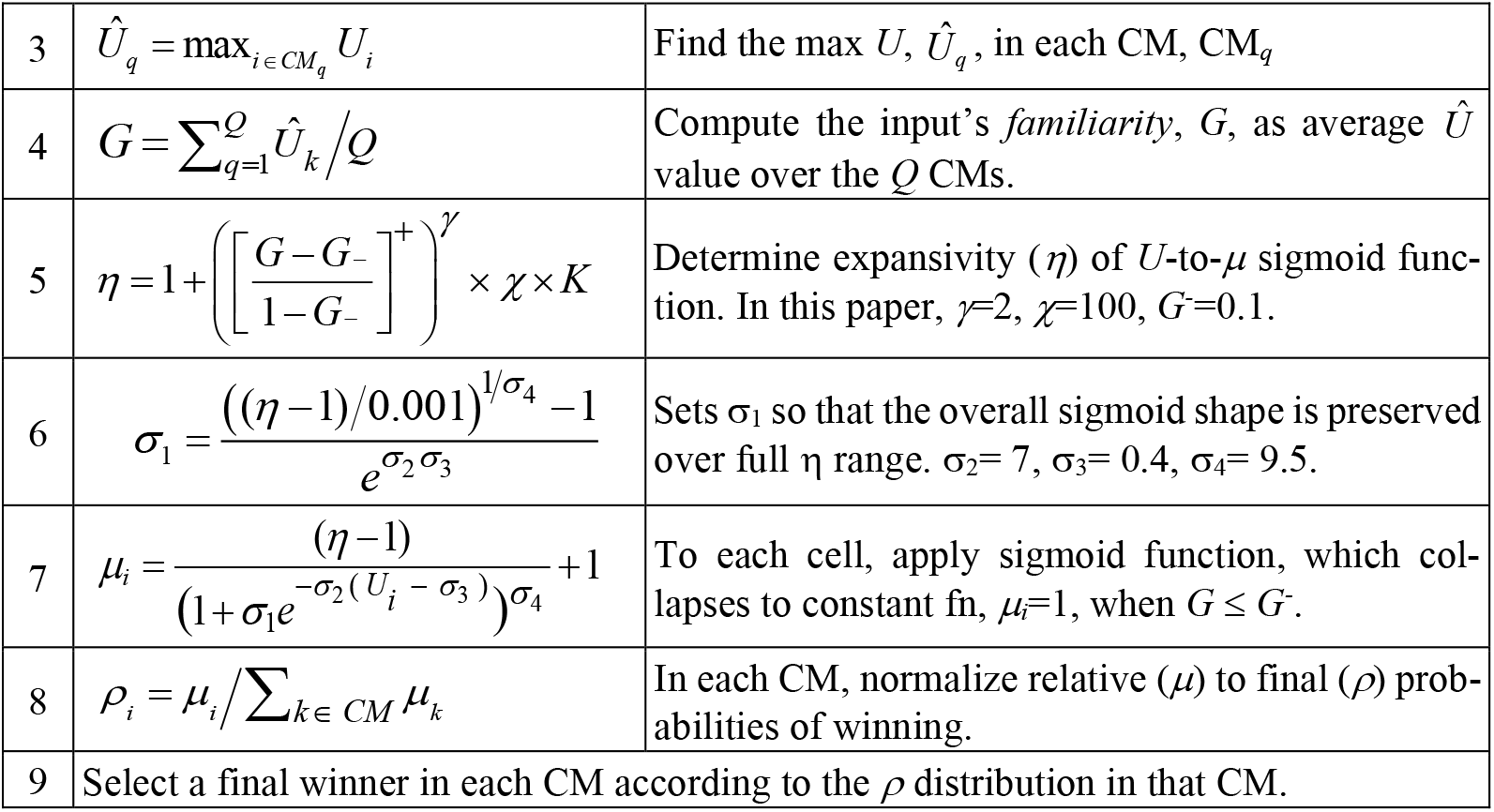
Simplified Code Selection Algorithm (CSA)

a. When high global familiarity is detected (*G*≈1), those distributions are exaggerated to bias the choice in favor of cells that have high input summations, and thus, high *local* familiarities (*U*), which acts to increase correlation.
b. When low global familiarity is detected (*G*≈0), those distributions are flattened so as to reduce bias due to local familiarity, which acts to increase the expected Hamming distance between the selected code and previously stored codes, i.e., to decrease correlation.

Since the *U* values represent *signal*, exaggerating the *U* distribution in a CM increases signal whereas flattening it increases noise. The above behavior (and its smooth interpolation over the range, *G=*1 to *G=*0) is the means by which Sparsey achieves SISC. And, it is the enforcement (statistically) of SISC during learning, which ultimately makes possible the immediate, i.e., *fixed time* (the number of algorithmic steps needed to do the retrieval is independent of the number of stored codes), retrieval of the best-matching (most likely, most relevant) hypothesis.

## 3 Results

The simulation-backed example of this section demonstrates that the CSA achieves the property, i.e., statistical (approximate) preservation of similarity from inputs to codes, qualitatively described in Fig. 2. In the experiment, the six inputs, *I*_1_ to *I*_6_, at top of Fig. 3a, were presented once each and assigned to the codes, *ϕ*(*I*_*1*_) to *ϕ*(*I*_*6*_) (not shown), via execution of the CSA (Table 1). The six inputs are disjoint only for simplicity of exposition. The input field (receptive field, RF) is a 12×12 binary pixel array and all inputs are of the same size, 12 active pixels. Since all inputs have exactly 12 active pixels, input similarity is simply *sim*(*I*_*x*_,*I*_*y*_) = |*I*_*x*_⋂*I*_*y*_|/12, shown as decimals under inputs. The CF consists of *Q=*24 WTA CMs, each having *K=*8 binary cells. The second row of Fig. 3a shows a novel stimulus, *I*_7_, and its varying overlaps (yellow pixels) with *I*_1_ to *I*_6_. Fig. 3b shows the code, *ϕ*(*I*_*7*_), activated (by the CSA) in response to presentation of *I*_7_. Black indicates cells that also won for *I*_1_, red indicates active cells that did not win for *I*_1_, and green indicates inactive cells that did win for *I*_1_. Fig. 3c shows (using the same color interpretations) the detailed values of all relevant variables (*u, U, μ*, and *ρ*) computed by the CSA when *I*_7_ presents, and the winners drawn from the *ρ* distribution in each of the *Q*=24 CMs.

**Fig. 3.**
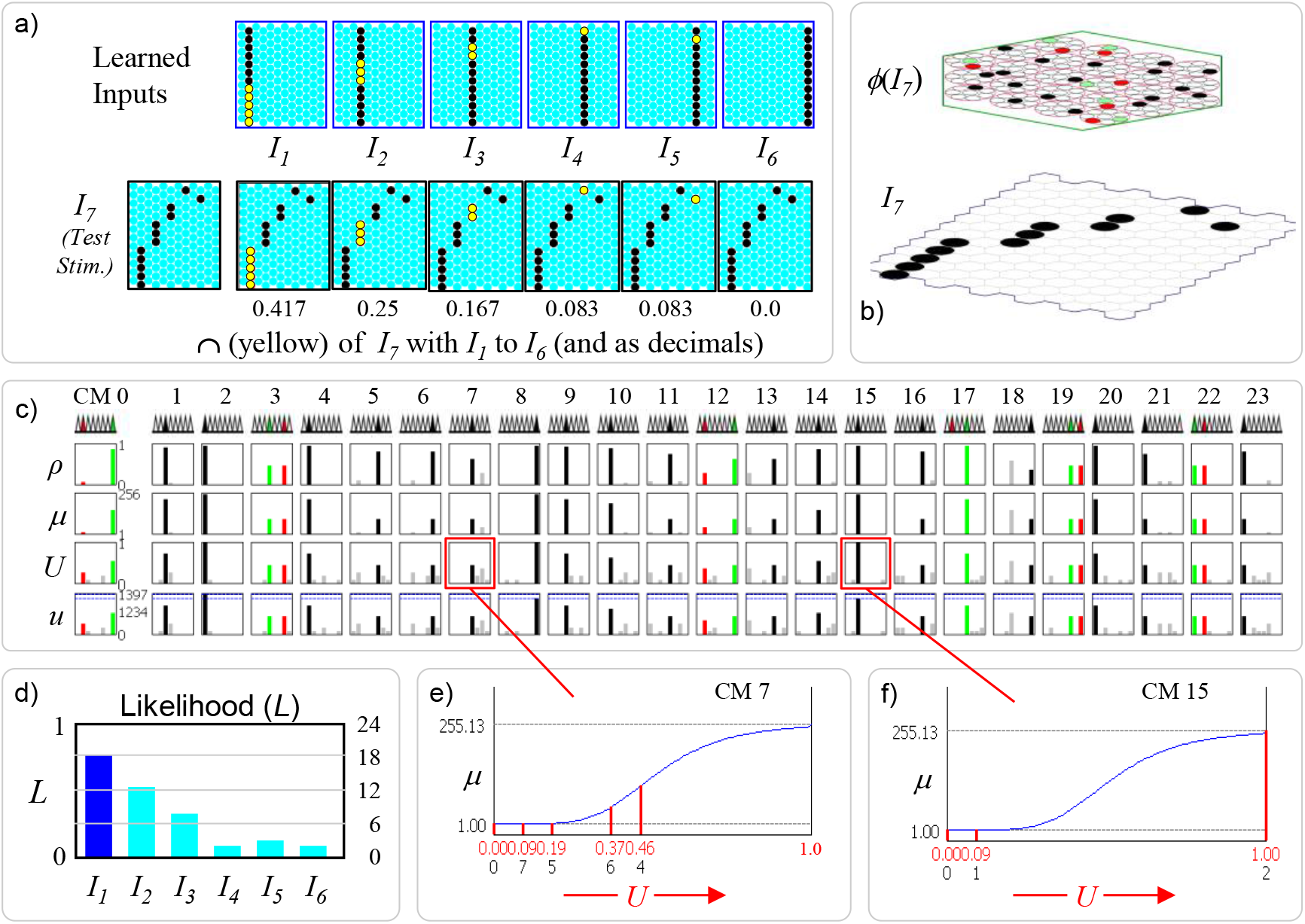
In response to a novel input, *I*_7_, the codes for the six previously learned (stored) inputs, *I*_1_ to *I*_6_, i.e., hypotheses, are activated with strength approximately correlated with the similarity (pixel overlap) of *I*_7_ input and those stored inputs. Test input *I*_7_ is most similar to learned input, *I*_1_, shown by the intersections (yellow pixels) in panel a. Thus, the code with the largest fraction of active cells is *ϕ*(*I*_*1*_) (18/24=75%) (blue bar in panel d). The other codes of the other inputs are active in rough proportion to their similarities with *I*_7_ (cyan bars). (c) Raw (*u*) and normalized (*U*) input summations to all cells in all CMs. Note: all weights are effectively binary, though “1” is represented with 127 and “0” with 0. Hence, the max *u* value possible in any cell when *I*_7_ is presented is 12×127=1524. The *U* values are transformed to un-normalized win probabilities (*μ*) in each CM via a sigmoid transform whose properties, e.g., max value of 255.13, depend on *G* and other parameters. *μ* values are normalized to true probabilities (*ρ*) and one winner is chosen in each CM (indicated in row of triangles: black: winner for *I*_7_ that also won for *I*_1_; red: winner for *I*_7_ that did not win *I*_1_: green: winner for *I*_1_ that did not win for *I*_7_. (e, f) Details for CMs, 7 and 15. Values in lower row of *U* axis are indexes of cells (within the CM) having the *U* values above them (red). Some CMs have a single cell with much higher *U* (and thus *ρ*) value than the rest (e.g., CM 15), some CMs have two cells tied for the max (CMs 3, 19, 22).

If we consider presentation of *I*_7_ to be a retrieval test, then the desired result is that the code of the most similar stored input, *I*_1_. should be retrieved (reactivated). In this case, the red and green cells in a given CM can be viewed as substitution errors, i.e., the green cell had the max *U* value in the CM and should have been reactivated, but since the final winner is a draw, occasionally a cell with a (possibly much) lower *U* value wins (CMs, 0, 12, 17). However, these are sub-symbolic scale errors, not errors at the scale of whole inputs (hypotheses), as a whole input is *collectively* represented by the entire MSDC code (entire *cell assembly*). In this example, appropriate threshold settings in downstream computations, would allow the model as a whole to return the correct answer given that 18 out of 24 cells of *I*_1_’s code, *ϕ*(*I*_*1*_), are activated, similar to thresholding schemes in other associative memory models [40, 41].

More generally, when *I*_7_ Is presented, we would like *all* of the stored inputs to be reactivated in proportion to their similarities to the test probe, *I*_7_. Fig. 3d shows that this approximately occurs. The active fractions of the codes, *ϕ*(*I*_*1*_) to *ϕ*(*I*_*6*_), are highly rank-correlated with the pixel-wise similarities of the corresponding inputs to *I*_7_. Thus, the blue bar in Fig. 3d represents the fact that the code, *ϕ*(*I*_*1*_), for the best matching stored input, *I*_1_, has the highest active code fraction, 75% (18 out 24, the black cells in Fig. 3b) cells of *ϕ*(*I*_*1*_) are active in *ϕ*(*I*_*7*_). The cyan bar for the next closest matching stored input, *I*_2_, indicates that 12 out of 24 of the cells of *ϕ*(*I*_*2*_) (code note shown) are active in *ϕ*(*I*_*7*_). In general, many of these 12 may be common to the 18 cells in {*ϕ*(*I*_*7*_)⋂*ϕ*(*I*_*1*_)}. And so on for the other stored hypotheses. [Note that even the code for *I*_6_ which has zero intersection with *I*_7_ has two cells in common with *ϕ*(*I*_*1*_). In general, the expected code intersection for the zero input intersection condition is not zero, but chance, since in that case, the winners are chosen from the uniform distribution in each CM, in which case the expected intersection is *Q/K*.]

If, instead of viewing presentation of *I*_7_ as a retreival test, we view it as a learning trial, we want the sizes of intersection of the code, *ϕ*(*I*_*7*_), activated in response, with the six previously stored codes, *ϕ*(*I*_*1*_) to *ϕ*(*I*_*6*_), to approximately correlate with the similarities of *I*_7_ to inputs, *I*_1_ to *I*_6_. But again, this is what Fig. 3d shows. As noted earlier, we assume that the similarity of a stored input *I*_x_ to the current input can be taken as a measure of *I*_x_’s probability/likelihood. And, since all codes are of size *Q*, we can divide code intersection size by *Q*, yielding a measure normalized to [0,1]: *L*(*I*_*1*_)=|*ϕ*(*I*_*7*_)⋂ *ϕ*(*I*_*1*_)|/Q. Thus, this result demonstrates that the CSA, a single-trial, unsupervised, non-optimization-based, and *most importantly, fixed time*, algorithm statistically enforces SISC. In this case, the red cells would not be considered errors: they would just part of a new code, *ϕ*(*I*_*7*_), being assigned to represent a novel input, *I*_7_, in a way that respects similarity in the input space. Crucially, because all codes are stored in *superposition* and because, when each one is stored, it is stored in a way that respects similarities with all previously stored codes, the patterns of intersection amongst the set of stored codes reflects not simply the *pairwise* similarity structure over the inputs, but, in principle, the similarity structure *of all orders present* in the input set. This is similar in spirit to another neural probabilistic model [2, 42], which proposes that overlaps of distributed codes (and recursively, overlaps of overlaps), encode the domain’s latent variables (their identities and valuednessess), cf. “anonymous latent variables”, [43].

The likelihoods in Fig. 3d may seem high. After all, *I*_7_ has less than half its pixels in common with *I*_1_, etc. Given these particular input patterns, is it really reasonable to consider *I*_1_ to have such a high likelihood? Bear in mind that our example assumes that the only experience this model has of the world are single instances of the six inputs shown. We assume no prior knowledge of any underlying statistical structure generating the inputs. Thus, it is really only the *relative* values that matter and we could pick other parameters, notably in CSA Steps 5-7, that would result in a much less expansive sigmoid nonlinearity, which would result in lower expected intersections of *ϕ*(*I*_*7*_) with the learned codes, and thus lower likelihoods. The main point is simply that the expected code intersections correlate with input similarity, and thus, likelihood.

A cell’s *U* value represents the total *local evidence* that it should be activated. However, rather than simply picking the max *U* cell in each CM as winner (i.e., hard max), which would amount to executing only steps 1-3 of the CSA, the remaining CSA steps, 4-9, are executed, in which the *U* distributions are transformed as described earlier and winners are chosen as draws (shown in the row of triangles just below CM indexes) from the *ρ* distributions in each CM. Thus, an extremely cheap-to-compute (CSA Step 4) *global* function of the whole CF, *G*, is used to influence the *local* decision process in each CM. We repeat for emphasis that no part of the CSA explicitly operates on, i.e., iterates over, stored hypotheses (codes); indeed, there are no explicit (localist) representations of stored hypotheses on which to operate.

Fig. 4 shows that presentation of different novel inputs yields different likelihood distributions that correlate approximately with similarity. Input *I*_8_ (Fig. 4a) has highest intersection with *I*_2_ and a different pattern of intersections with the other learned inputs as well (refer to Fig. 3a). Fig. 4c shows that the codes of the stored inputs become active in approximate proportion to their similarities with *I*_8_, i.e., their likelihoods are simultaneously physically represented by the fractions of their codes which are active. The *G* value in this case, 0.65, yields, via CSA steps 5-7, the *U*-to-*μ* transform shown in Fig. 4b, which is applied in all CMs. Its range is [1,300] and given the particular *U* distributions shown in Fig. 4d, the cell with the max *U* in each CM ends up being greatly favored over other lower-*U* cells. The red box shows the *U* distribution for CM 9. The second row of the abscissa in Fig. 4b gives the within-CM indexes of the cells having the corresponding (red) values immediately above (shown for only three cells). Thus, cell 3 has *U=*0.74 which maps to approximately *μ* ≈ 250 whereas its closest competitors, cells 4 and 6 (gray bars in red box) have *U=*0.19 which maps to *μ* = 1. Similar statistical conditions exist in most of the other CMs. However, in three of them, CMs 0, 10, and 14, there are two cells tied for max *U*. In two, CMs 10 and 14, the cell that is not contained in *I*_2_’s code, *ϕ*(*I*_*2*_), wins (red triangle and bars), and in CM 0, the cell that is in *ϕ*(*I*_*2*_) does win (black triangle and bars). Overall, presentation of *I*_8_ activates a code *ϕ*(*I*_*8*_) that has 21 out of 24 cells in common with *ϕ*(*I*_*2*_) manifesting the high likelihood estimate for *I*_2_.

**Fig. 4.**
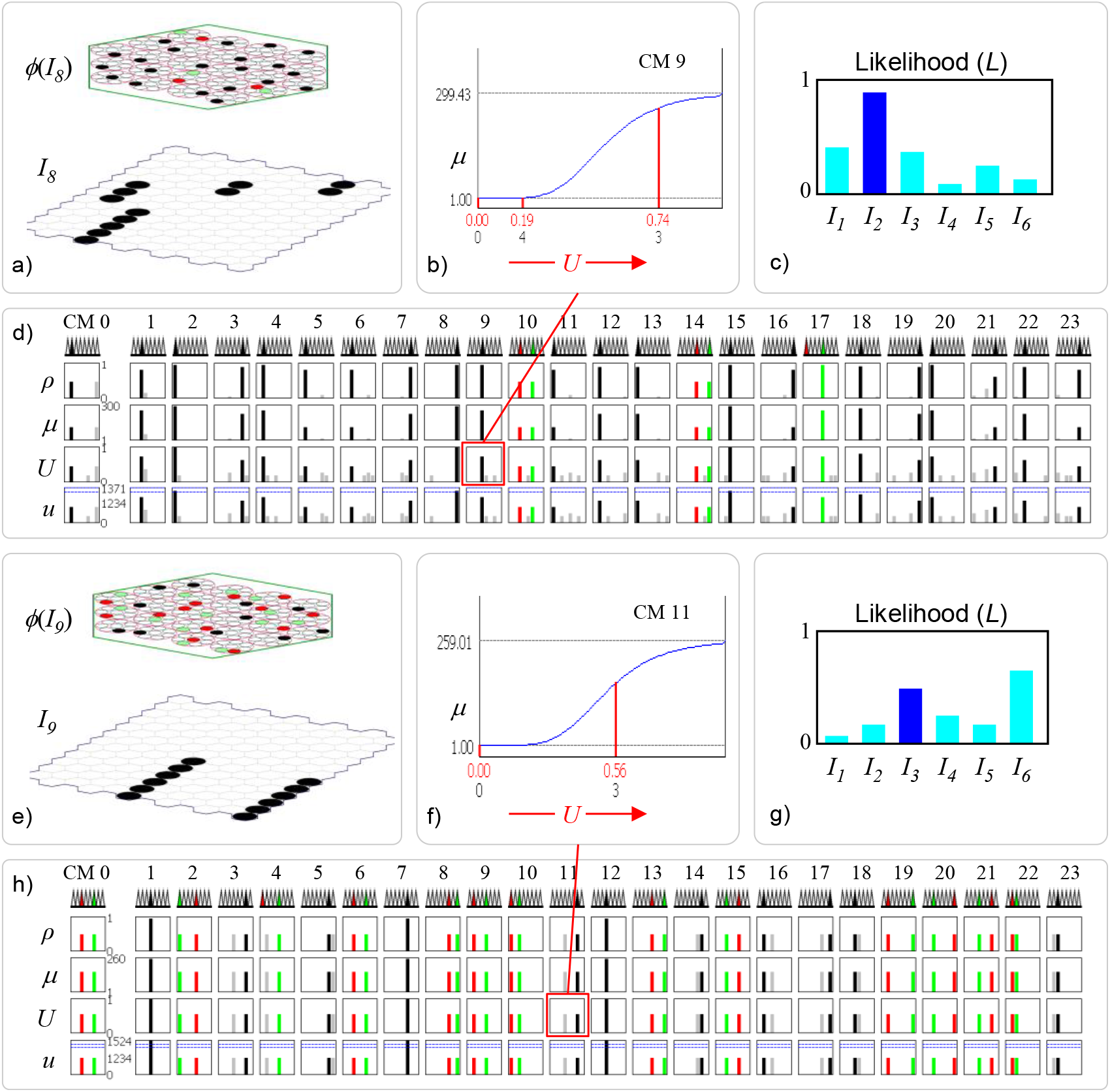
Details of presenting other novel inputs, *I*_8_ (panels a-d) and *I*_9_ (panels e-h). In both cases, the resulting likelihood distributions (panels c,g) correlate closely with the input overlap patterns. Panels b and f show details of one example CM (red boxes in panels d and h) for each input.

Finally. Fig. 4e shows presentation of a more ambiguous input, *I*_9_, having half its pixels in common with *I*_3_ and the other half with *I*_6_. Fig. 4g shows that the codes for *I*_3_ and *I*_6_ have both become approximately equally (with some statistical variance) active and are both more active than any of the other codes. Thus, the model is representing that these two hypotheses are the most likely and approximately equally likely. The exact bar heights fluctuate somewhat across trials, e.g., sometimes *I*_3_ has higher likeli-hood than *I*_6_, but the general shape of the distribution is preserved. The remaining hypotheses’ likelihoods also approximately correlate with their pixelwise intersections with *I*_9_. The qualitative difference between presenting *I*_8_ and *I*_9_ is readily seen by comparing the *U* rows of Fig. 4d and 4h and seeing that for the latter, a tied max *U* condition exists in almost all the CMs, reflecting the equal similarity of *I*_9_ with *I*_3_ and *I*_6_. In approximately half of these CMs the cell that wins intersects with *ϕ*(*I*_*3*_) and in the other half, the winner intersects with *ϕ*(*I*_*6*_). In Fig. 4h, the three CMs in which there is a single black bar, CMs 1, 7, and 12, indicates that the codes, *ϕ*(*I*_*3*_) and *ϕ*(*I*_*6*_), intersect in these three CMs.

### 3.1 A MSDC simultaneously transmits the full likelihood distribution via an atemporal combinatorial spike code

The use of MSDC allows the likelihoods of *all* hypotheses stored in the distribution, to be transmitted via a set of simultaneous single spikes from the neurons comprising the active MSDC. This is shown in the example given in Fig. 5e, which, at the same time, compares this fundamentally new *atemporal, combinatorial spike code*, with temporal spike codes and one prior (in principle) atemporal code. For a single source neuron, two types of spike code are possible, rate (frequency) (Fig. 5b), and latency (e.g., of spike(s) relative to an event, e.g., phase of gamma) (Fig. 5c). Both are fundamentally temporal and have the crucial limitation that only one value (item) represented by the source neuron can be sent at a time. Most prior population-based codes also remain fundamentally temporal: the signal depends on spike *rates* of the afferent axons, e.g., [1-4] (not shown in Fig. 5).

**Fig. 5.**
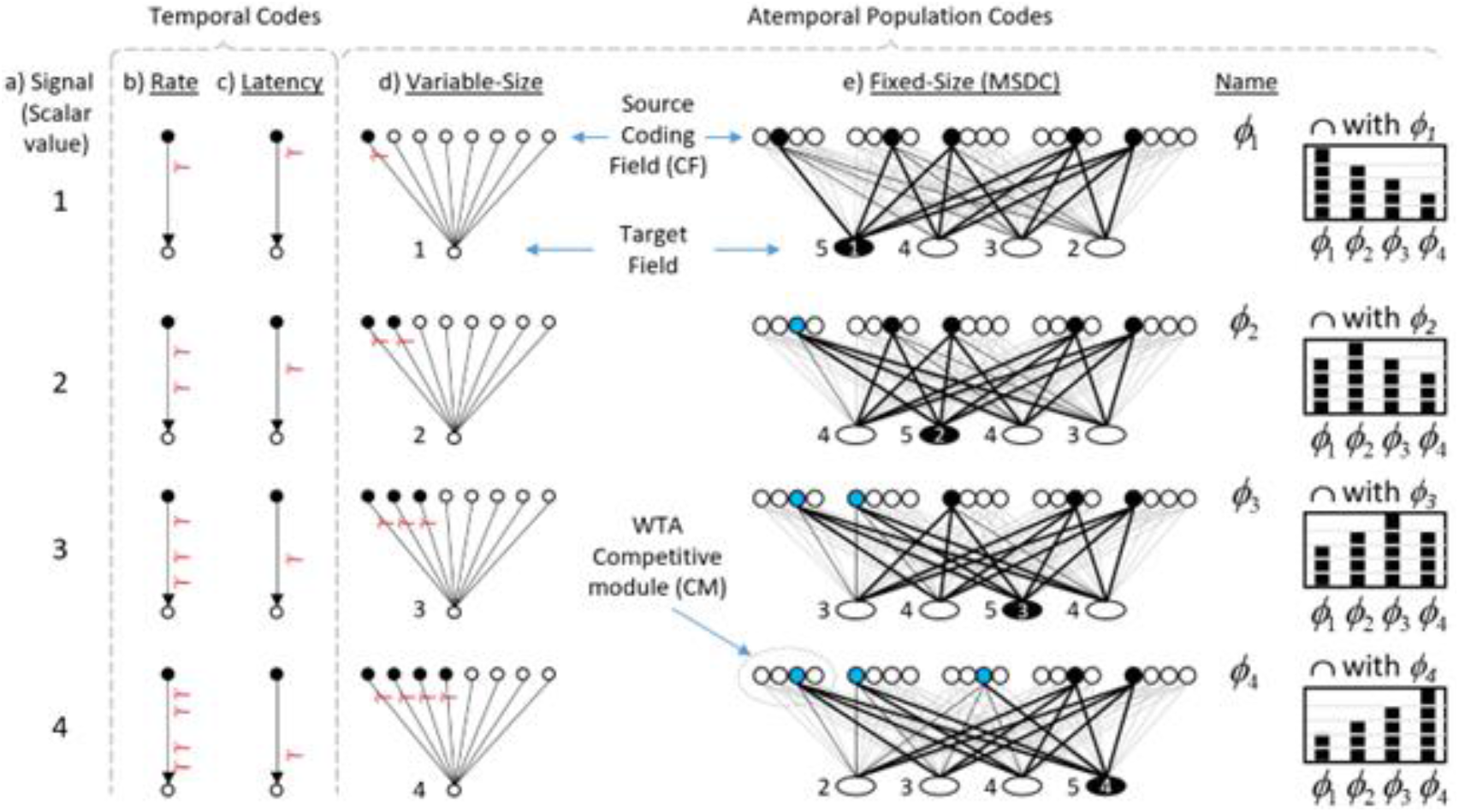
Temporal vs. atemporal spike coding concepts. The fixed-size MSDC code has the advantage of being able to send the entire distribution, i.e., the likelihoods of *all* codes (hypotheses) stored in the source CF, with a set of simultaneous single spikes from Q=5 units comprising an active MSDC code. See text for details.

Fig. 5d illustrates an (effectively) atemporal population code [5] in which the *fraction* of active neurons in a source field carries the message, coded as the number of simultaneously arriving spikes to a target neuron (shown next to the target neuron for each of the four signals values). This *variable-size* population (a.k.a. “thermometer”) code has the benefit that all signals are sent in the same, short time, but it is not combinatorial in nature, and has limitations, including: a) the max number of representable values (items/concepts) is the number (*N*) of units comprising the source CF; and b) as for the temporal codes defined with respect to a single source neuron, any single message sent can represent *only one* item, e.g., a single value of a scalar variable, i.e., implying that any one message carries only log_2_*N* bits.

In contrast, consider the *fixed-size* MSDC code of Fig. 5e. The source CF consists of *Q=*5 CMs, each with *K=*4 binary units. Thus, all codes, are of the same fixed size, *Q*=5. As done in Fig. 2, the codes for this example were manually chosen to reflect the similarity structure of scalar values (Col. a) (the prior section has already demonstrated that the CSA statistically preserves similarity). As suggested by charts at right of Fig. 5, any single MSDC, *ϕ*_i_, represents (encodes) the similarity distribution over *all* items (values) stored in the field. Note: blue denotes active units *not* in the intersection with *ϕ*_1_. We’re assuming that input (e.g., scalar value) similarity correlates with likelihood, which again, is reasonable for vast portions of input spaces having natural statistics.

Since any one MSDC, *ϕ*_i_, encodes the full likelihood distribution, the set of single spikes sent from it simultaneously transmits that full distribution, encoded as the instantaneous sums at the target cells. Note: when any MSDC, *ϕ*_i_, is active, 20 wts (axons) will be active (black), thus, all four target cells will have *Q*=5 active inputs. Thus, due to the *combinatorial* nature of the MSDC code, the specific values of the binary weights are essential to describing the code (unlike the other codes where we can assume all wts are 1). Thus, for the example of Fig. 5e, we assume: a) all wts are initially 0; b) the four associations, *ϕ*_1_ → target cell 1, *ϕ*_2_ → target cell 2, etc., were previously stored (learned) with single trials; and c) on those learning trials, coactive pre-post synapses were increased to *w*=1. Thus, if *ϕ*_1_ is reactivated, target cell 1’s input sum will be 5 and other cells’ sums will be as shown (to left of target cells). If *ϕ*_2_ is reactivated, target cell 2’s input sum will be 5, etc. [Black line: active *w*=1; dotted line: active *w*=0; gray line: *w*=0.] As described in Fig. 3 of [7], the four target cells could be embedded in a recurrent field with inhibitory infrastructure allowing sequential read out in descending input sum order, implying that the full similarity (likelihood) order information over all four stored items is sent in each of the four cases. As there are 4! orderings of the four items, each such message, each a set of 20 simultaneous spikes sent from five active CF units, sends log_2_(4!)=4.58 bits. We suggest this marriage of fixed-size MSDCs and an atemporal spike code is a crucial advance beyond prior population-based models, i.e., the “distributional encoding” models (see [8, 9] for reviews), and may be key to explaining the speed and efficiency of probabilistic computation in the brain.

## 4 Discussion

We described a radically different theory, from prevailing probabilistic population coding (PPC) theories, for how the brain represents and computes with probabilities. This theory, Sparsey, avails itself only in the context of *modular sparse distributed coding* (MSDC), as opposed to the fully distributed coding context in which the PPC models have been developed (or a localist context). The theory, Sparsey, was introduced 25+ years ago, as a model of the canonical cortical circuit and a computationally efficient explanation of episodic and semantic memory for sequences, but its interpretation as a way of representing and computing with probabilities was not emphasized. The PPC models [5, 7-10, 12, 42] share several fundamental properties: 1) continuous neurons; 2) full/dense coding; 3) due to 1 and 2, synapses must either be continuous or rate coding must be used to allow decoding; 4) they generally assume rate coding; 5) individual neurons are generally assumed to have unimodal, e.g., bell-shaped, tuning functions (TFs); 6) individual neurons are assumed to be noisy, and noise is generally viewed as degrading computation, thus, needing to be mitigated, e.g., averaged out.

In contrast to these PPC properties/assumptions, Sparsey assumes: 1) binary neurons; 2) items of information are represented by small (relative to whole CF) sets of neurons (MSDCs) and any such code simultaneously represents not only the likelihood of the single best matching stored hypothesis, but, simultaneously, the likelihoods of *all* stored hypotheses; 3) only effectively binary synapses; 4) signaling via waves of simultaneous single (e.g., first) spikes from a source MSDC; 5) all weights are initially zero, i.e., the TFs are initially completely flat, and emerge via single/few-trial, unsupervised learning to reflect a neuron’s specific history of inclusion in MSDCs; 6) rather than being viewed as a problem imposed by externalities (e.g., common input, intrinsically noisy cell firing), noise functions as a resource, controlled usage of which yields the valuable property that similar inputs are mapped to similar codes (SISC).

The CSA’s algorithmic efficiency, i.e., both learning (storage) and best-match retrieval are fixed time operations, has not been shown for any other computational method, including hashing methods, either neurally-relevant [44-46], or more generally [reviewed in [47]]. Although time complexity considerations like these have generally not been discussed in the PPC literature, they are essential for evaluating the overall plausibility of models of biological cognition, for while it is uncontentious that the brain computes probabilistically, we also need to explain the extreme speed with which these computations, over potentially quite large hypothesis spaces, occur.

One key to Sparsey’s computational speed is its extremely efficient method of computing the *global* familiarity, *G*, simply as the average of the max *U* values of the *Q* CMS. In particular, computing *G does not require* explicitly comparing the new input to every stored input (nor to a log number of the stored inputs as is the case for tree-based methods). *G* is then used to adjust, in the same way, the transfer functions of all neurons in a CF. This dynamic, and fast timescale (e.g., 10 ms), modulation of the transfer function, based on the *local* (to the CF, thus *mesoscale* circuit) measure, *G*, is a strongly distinguishing property of Sparsey: in most models, the transfer function is static. While there has been much discussion about the nature, causes, and uses, of correlations and noise in cortical activity; see [48-50] for reviews, the *G*-based titration of the amount of noise present in the code selection process, to achieve the specific goal of approximately preserving similarity (SISC) is a novel contribution to the discussion.

Enforcing SISC in the context of an MSDC CF realizes a balance between:

a. maximizing the storage capacity of the CF, and
b. embedding the similarity structure of the input space in the set of stored codes, which in turn enables fixed-time best-match retrieval.

In exploring the implications of shifting focus from information theory to coding theory viz. theoretical neuroscience, [51] pointed to this same tradeoff, though their treatment uses error rate (coding accuracy) instead of storage capacity. Understanding how neural correlation ultimately affects things like storage capacity is considered largely unknown and an active area of research [52]. Our approach implies a straightforward answer. Minimizing correlation, i.e., maximizing average Hamming distance over the set of codes stored in an MSDC CF, maximizes storage capacity. Increases of any correlations of pairs, triples, or subsets of any order, of the CF’s units increases the strength of embedding of statistical (similarity) relations in the input space.

Sparsey has many more features and capabilities than can be described here, e.g., it has been generalized to the temporal domain and to hierarchies of CFs. Nevertheless, the results shown here will hopefully pique further interest.

